# Meningitis pathogens evade immune responses by thermosensing

**DOI:** 10.1101/586131

**Authors:** Hannes Eichner, Laura Spelmink, Anuj Pathak, Birgitta Henriques-Normark, Edmund Loh

**Affiliations:** Department of Microbiology, Tumor and Cell Biology, Karolinska Institutet, 171 77 Stockholm, Sweden; Department of Clinical Microbiology, Karolinska University Hospital, 171 65 Stockholm, Sweden; SCELSE and LKC, Nanyang Technological University, 639798 Singapore

## Abstract

Bacterial meningitis is a major cause of death and disability in children worldwide. Two human restricted pathogens, *Streptococcus pneumoniae* and *Haemophilus influenzae*, are the major causative agents of bacterial meningitis, attributing to 200,000 deaths annually. These pathogens are often part of the nasopharyngeal microflora of healthy carriers. However, what factors elicit them to disseminate and cause invasive diseases remain unknown. Elevated temperature and fever are hallmarks of inflammation triggered by infections and can act as warning signal to these pathogens. Here, we investigate whether these pathogens could sense environmental temperature to evade host complement-mediated killing. We show that expression of two vital virulence factors and vaccine components, the capsule and factor H binding proteins, are temperature dependent. We identify and characterize four novel RNA thermosensors in *S. pneumoniae* and *H. influenzae* within their 5′-untranslated regions of genes, responsible for capsular biosynthesis and production of factor H binding proteins. Our data further demonstrate that these pathogens have co-evolved thermosensing abilities independently with unique RNA sequences, but distinct secondary structures, to evade the human immune system.

**Author Summary:** *Streptococcus pneumoniae* and *Haemophilus influenzae* are bacteria that reside in the upper respiratory tract. This harmless colonization may progress to severe and often lethal septicaemia and meningitis, but molecular mechanisms that control why these pathogens invade the circulatory system remain largely unknown. Here we show that both *S. pneumoniae* and *H. influenzae* can evade complement killing by sensing the temperature of the host. We identify and characterize four novel RNA thermosensors in *S. pneumoniae* and *H. influenzae* within their respective 5′-untranslated regions of genes, influencing capsular biosynthesis and production of factor H binding proteins. Moreover, we show that these RNA thermosensors evolved independently with exclusive unique RNA sequences to sense the temperature in the nasopharynx and in other body sites to avoid immune killing. Our finding that regulatory RNA senses temperatures and directly regulate expression of two important virulence factors and vaccine components of *S. pneumoniae* and *H. influenzae*, is most important for our understanding of bacterial pathogenesis and for vaccine development. Our work could pave the way for similar studies in other important bacterial pathogens and enables clinicians and microbiologists to adjust their diagnostic techniques, and treatments to best fit the condition of the patients.

## Introduction

*Streptococcus pneumoniae* and *Haemophilus influenzae* are both human restricted pathogens that may cause deadly meningitis and sepsis. In order to survive in the host, these bacteria have evolved analogous survival mechanisms such as the production of polysaccharide capsules to evade host immune responses. Encapsulated bacterial pathogens pose a major threat to human health and contribute to significant morbidity and mortality[1]. For Gram-positive pneumococci, the capsule enables the bacteria to avoid phagocytosis, and at least 97 different capsular serotypes have been identified as of 2015[2]. Pneumococci have been suggested to be resistant to complement killing due to their thick layer of peptidoglycan[3]. Gram-negative *H. influenzae* shows six capsular serotypes, facilitating the bacterium to resist phagocytosis and complement-mediated killing[4]. Vaccines based on bacterial polysaccharide capsules are being used to prevent *S. pneumoniae* and *H. influenzae* infections[5].

In addition to expression of capsules, these two pathogens also evade complement-mediated killing by binding to human Factor H (FH), the major negative regulator of the alternative complement pathway. FH is recruited by high affinity interactions with FH binding proteins expressed on the bacterial surface, such as Pneumococcal Surface Protein C (PspC) and the Protein H (PH, encoded by the *lph* gene) for *H. influenzae*[6-8]. FH binding proteins have been included in a vaccine against *Neisseria meningitidis*, but have also been suggested as vaccine candidates against pneumococcal infections[9-11].

The ecological niche for *S. pneumoniae* and *H. influenzae* is the nasopharyngeal cavity from where they may spread into the bloodstream and cause disease. There is a temperature gradient in the nasal cavity where these two pathogens colonize. The temperature on the surface of the anterior nares is around 30°C to 32°C at the end of inspiration, and rises to around 34°C in the posterior nasopharynx and tonsillar region[12, 13]. Both these sites on the mucosal surface are significantly cooler than the core body temperature of 37°C, where the bacteria replicate during invasive diseases. Previously, we showed that temperature acts as a danger signal to *N. meningitidis*, prompting the bacterium to enhance expression of immune evasion factors via three independent RNA thermosensors[14]. RNA thermosensors are elements usually located in the 5′-untranslated region (5′-UTR) of an mRNA transcript, forming a secondary structure at lower temperatures that inhibits protein translation by blocking access of ribosomes to the ribosome binding site (RBS). As temperature rises, the RNA secondary structure undergoes a conformational change due to higher thermodynamic energy, exposing the RBS, and thus allowing translation[15].

The importance of temperature sensing in meningococci prompted us to investigate whether temperature affects expression of virulence factors in other meningitis causing pathogens that colonize the same niche, i.e. *S. pneumoniae* and *H. influenzae.* Although the role of the capsule and FH binding proteins in *S. pneumoniae* and *H. influenzae* in complement evasion has been studied, their regulatory mechanisms remain largely unknown. Here we identify that thermosensing governs the expression of the capsular polysaccharide and FH binding proteins in *S. pneumoniae* and *H. influenzae*, thereby influencing complement-mediated evasion. Moreover, through sequence analyses of their 5′-UTRs, we found that the three meningitis causing pathogens have independently evolved unique RNA sequences, with no sequence conservation, to serve the same thermosensing function.

## Results

The pneumococcal capsular polysaccharide is encoded by a cluster of 10-20 tightly-linked genes[16]. The first four genes, that are located at the 5′-end of the capsular locus (*cpsABCD*), are common to all serotypes and are involved in the capsular regulation[17]. To investigate whether temperature is involved in capsular gene expression, we examined the expression of capsular polysaccharide synthesis protein B (CpsB) in the TIGR4 strain (of serotype 4)[18]. The TIGR4 strain was grown at three temperatures: 30°C, 37°C and 42°C with no observable growth defect. The expression of the CpsB protein from total protein lysates was consistently higher at elevated temperatures in three independent experiments (Fig 1A). As CpsA is co-transcribed and translated with CpsB, subsequent experiments were performed using CpsA. Two other pneumococcal proteins, not affected by temperatures, were used as controls (the autolysin LytA, and GAPDH (Glyceraldehyde-3-phosphate dehydrogenase) (Fig 1A).

**Figure 1.**
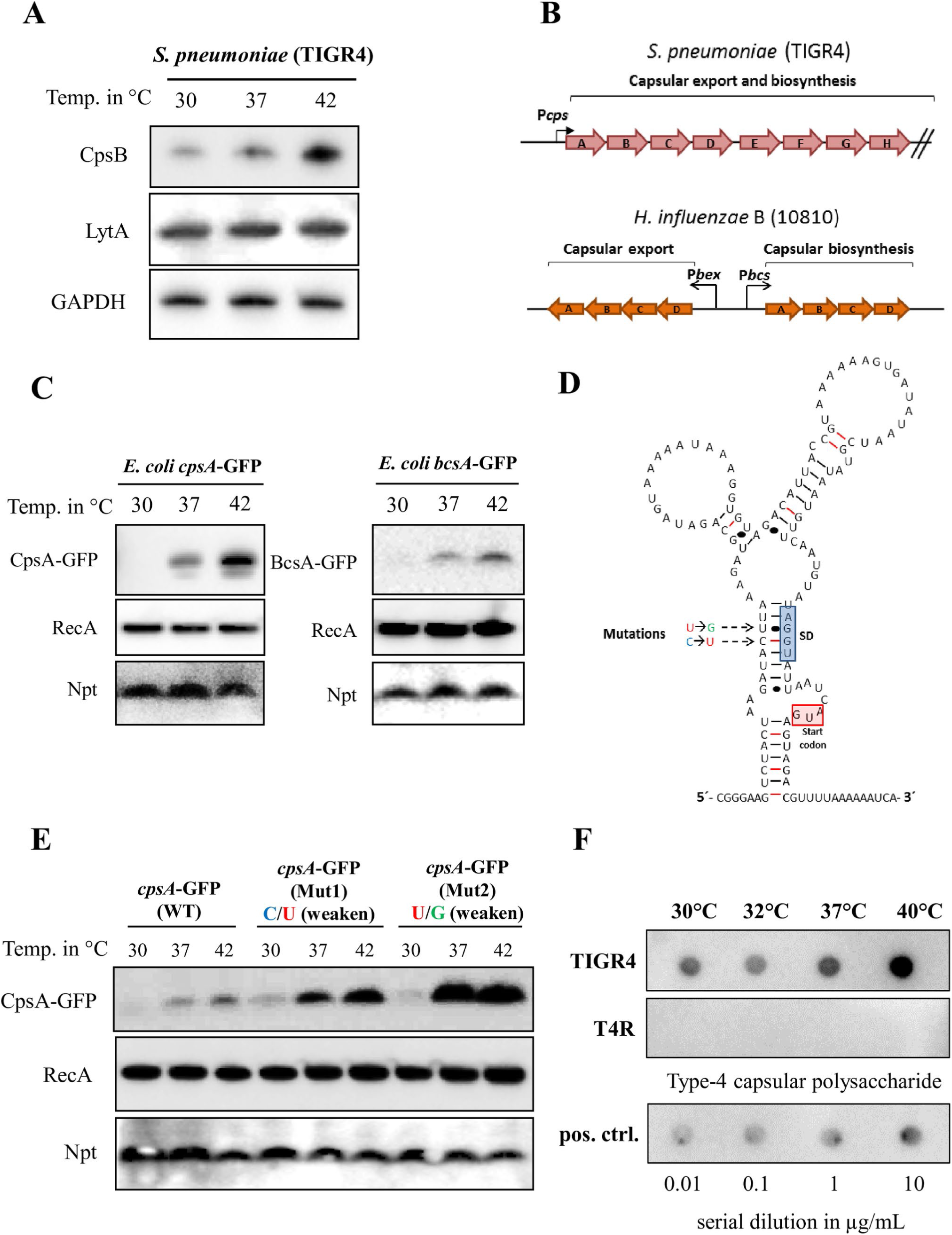
Capsular gene expression of S. pneumoniae and H. influenzae is temperature regulated. A) Western blot analysis of S. pneumoniae CpsB expression at different temperatures (LytA and GAPDH are used as loading controls). B) The capsular biosynthesis and transport operon map of S. pneumoniae (cps) and H. influenzae (bcs) with promoters (arrows). C) Thermoregulation of Cps and Bcs is detected in E. coli by western blot analysis (RecA and Neomycin-phosphotransferase are used as controls). D) Putative secondary structure of the cps 5′-UTR. The ribosome binding site (RBS) and start codon are indicated. E) Mutagenesis of the cps 5′-UTR. Western blot of E. coli expressing Cps mutations (position indicated in D) at different temperatures. F) Dot blots of S. pneumoniae show increment of capsular production with increasing temperatures. The non-encapsulated T4R strain is used as a negative control. Serial dilution of purified type-4 capsular polysaccharide (µg/mL).

Next, we examined the intergenic region sequences upstream of the capsular biosynthesis genes in both *S. pneumoniae* and *H. influenzae* for the presence of putative RNA thermosensors. By reviewing *S. pneumoniae* TIGR4 intergenic regions upstream of the capsular biosynthesis genes (Fig 1B) using the promoter prediction programme BPROM (Softberry Inc., Mt. Kisco, NY), we identified an additional novel putative σ-^70^ promoter, 146 base pairs (bps) upstream of the *S. pneumoniae cpsA* operon (S1A Fig). Similarly to *S. pneumoniae*, the capsular locus of *H. influenzae* comprises clustered regions encoding functions for capsular polysaccharide synthesis, modification, and translocation (Fig 1B)[19]. By investigating the serotype-specific region containing genes for capsular synthesis (*bcsABCD*) of *H. influenzae* type B, we identified a putative σ-^70^ promoter 215 bps upstream of the *bcsA* operon (S1B Fig).

RNA thermosensors are elements located within specific 5′-UTR of mRNAs and they could function in a heterologous host[14, 20]. Therefore, the whole intergenic regions upstream of *cpsA* and *bcsA* together with their respective first 24 coding-bases were translationally fused to a green fluorescent protein within the plasmid pEGFP-N2 and transformed into *Escherichia coli*. Thermoregulation of CpsA-GFP and BcsA-GFP was evident also in *E. coli* (Fig 1C). RecA antibody (Abcam) was used as a cytoplasmic loading control and anti-Neomycin-Phosphotransferase antibody (NovusBio) to eliminate potential effects of temperature on plasmid replication (Fig 1C). The two previously identified RNA thermosensors, the neisserial capsular CssA and the listerial transcriptional activator PrfA, were used as positive controls (S1C Fig). In addition, site-directed mutageneses were formed by inactivating either promoter-1 (Prom-1 Mut) or promoter 2 (Prom-2 Mut) by replacing the −35 sequences to investigate whether the longer or the shorter UTR is involved in thermosensing of CpsA (S2A Fig). Western blot analysis showed that both promoters of pneumococcal *cps* are required for production of Cps, however, only the longer transcript regulated by the novel promoter (promoter-1) is required for thermosensing (S2 Fig).

To eliminate temperature dependent transcription, the expression of *cpsA* and *bcsA* mRNAs was examined by Northern blotting. Both mRNA transcript levels were unaffected by temperature (S3 Fig), indicating that upregulation of CpsA and BcsA occurs post-transcriptionally. To further eliminate the involvement of host dependent factors in their thermoregulation, we subjected these GFP-fusion constructs to an *in vitro* transcription/translation assay at different biological relevant temperatures ranging from 30°C to 40^°^C (S4 Fig). Both CpsA and BcsA thermosensors displayed gradual increment of expression, similar to the rheostat expression of the meningococcal CssA[14]. In contrast, the PrfA thermosensor of the enteric pathogen *L. monocytogenes* displayed an on-off expression switch at 36-38°C (S4 Fig). Together, these results confirm the presence of *bona fide* RNA thermosensors controlling the capsular biosynthesis operons in *S. pneumoniae* and *H. influenzae.*

Using Vienna *RNAfold* package[21] and *VARNA* applet[22], we determined the putative secondary structure of *cpsA* and *bcsA* 5′-UTRs together with their first 24-coding bases. Consistent with the classical feature of a RNA thermosensor, the *cpsA* and *bcsA* 5′-untranslated region (UTR) regions form stem-loop structures that occlude the RBS (S5 Fig). Using the pneumococcal CpsA construct, we investigated the RNA thermosensor functionality by introducing nucleotide changes (+19_C/U_ and +20_U/G_) into the pneumococcal *cpsA* 5′-UTR predicted to alter the stability of the thermosensor by weakening the RBS base-pairing (Fig 1D). Expression of CpsA-GFP from the plasmids containing these changes was consistent with a thermosensor in the 5′-UTR (Fig 1E).

An elevated expression of the CpsA/CpsB proteins is however not a definite evidence that more capsule is produced. To address this, the production of capsular polysaccharides of *S. pneumoniae* was analyzed using a dot blot assay, as it has previously been used to quantify pneumococcal capsule production[23]. Dot blots using serotype 4 specific anti-sera showed that the polysaccharide capsule was produced in higher amounts with increasing temperature in the bacterial supernatant (Fig 1F).

To investigate whether other pneumococcal capsules could regulate the expression of their capsular genes by sensing temperature, three pneumococcal strains of important serotypes, 1, 2, and 22F, were studied. Thermoregulation of CpsB was evident also in these strains (S6A Fig) and their sequences of the *cpsA* 5’-UTR, important for forming the RNA secondary structure, were found to be conserved (S6B Fig). The conservation of the wild-type sequence and prevalence of an intact RNA-loop in these strains emphasize the importance of capsular thermoregulation in different pneumococcal isolates (S6C-E Figs).

We then examined whether other factors involved in immune evasion are subjected to similar thermoregulation and focused on FH binding proteins. Using an antibody recognizing the FH binding pneumococcal protein PspC, we observed that expression of PspC is also temperature dependent in different pneumococcal strains (Fig 2A and S6A Fig). Likewise, we also found that expression of the *H. influenzae* FH binding protein PH and its ability to bind to FH is also temperature dependent (Fig 2B) Expression of *pspC* and *lph* mRNA transcripts was unaffected by temperature changes (S3 Fig), suggesting that thermoregulation occurs post-transcriptionally, similar to their capsular gene counterparts. This finding prompted us to investigate the presence of RNA thermosensor structures within the 5′-region of pneumococcal *pspC* and the *H. influenzae lph* mRNAs. We reviewed the *pspC* and *lph* operons by using promoter prediction programs and identified two σ-^70^ promoters, 65 and 64 bps upstream of *S. pneumoniae pspC* and *H. influenzae lph* operons, respectively (S7A and S7B Figs).

**Figure 2.**
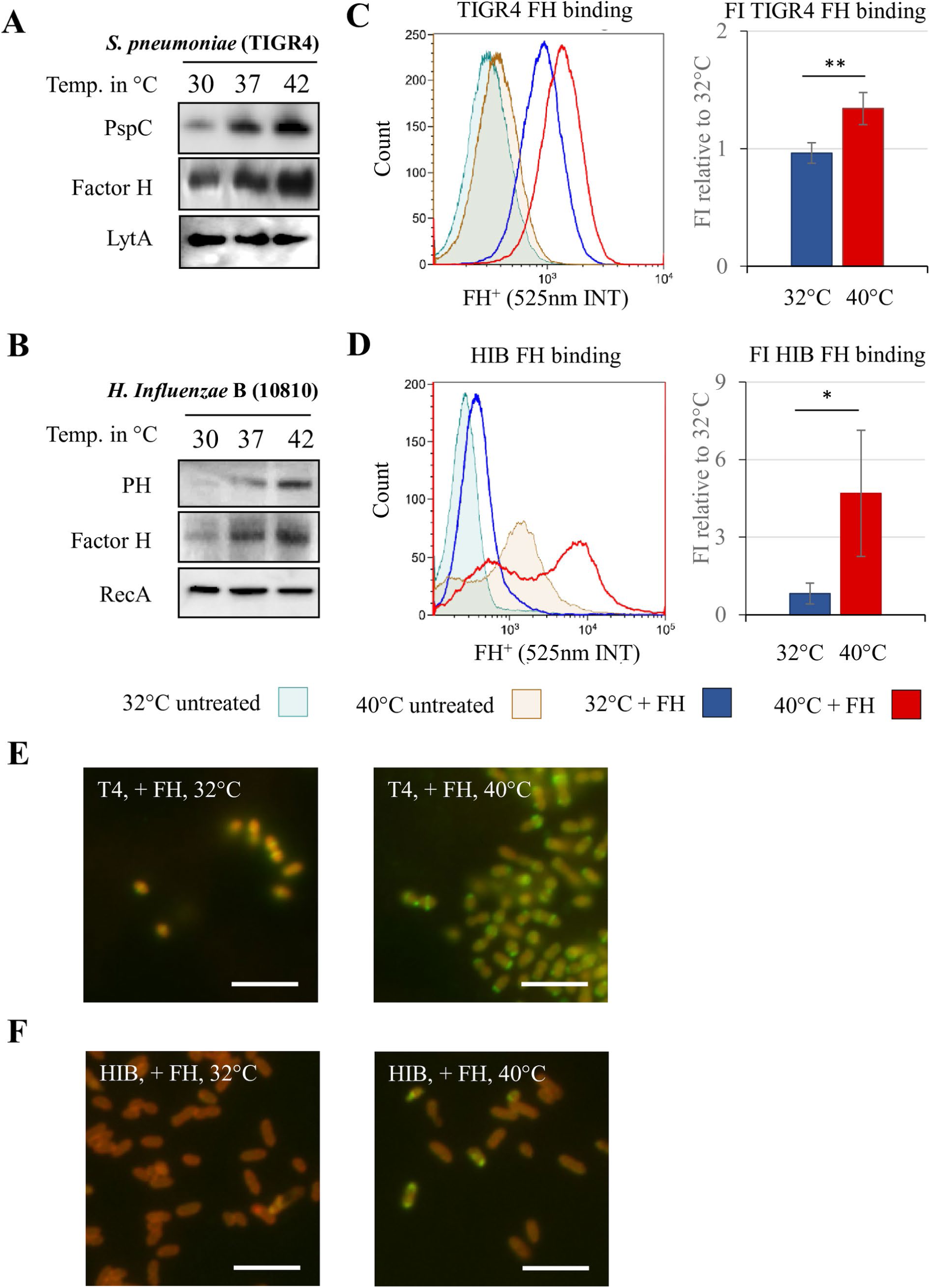
Factor H (FH) binding protein expression of *S. pneumoniae* and *H. influenzae* are temperature regulated. A) Western and far-western blot analyses show thermoregulated PspC and FH expression in *S. pneumoniae*. The autolysin LytA is used as loading control. B)Western and far-western blot analyses show thermoregulated PH expression and function in *H. influenzae*. RecA is used as loading control. C) Fluorescent flow cytometry shows increased human FH binding to *S. pneumoniae* when grown at higher temperatures (histograph). Fluorescence intensity of three experiments was pooled and FH binding of *S. pneumoniae* grown at 40°C compared to 32°C is shown (bar graph). D) Fluorescent flow cytometry shows two populations of *H. influenzae* with increased FH binding when grown at 40°C (histograph). E) Fluorescent microscopy analysis reveals that bacteria bind more FH at 40°C (more banded pattern of FH (green) binding to *S. pneumoniae* (auto-fluorescence of bacteria in red)) than at 32°C. Bar = 10µm.F) Fluorescent microscopy analysis reveals a FH (green) positive population of *H. influenzae* (auto-fluorescence in red) along with a FH negative population at 40°C. Bar = 10µm. Error bars denote s.e.m. * *p* < 0.05, ** p < 0.01 (Student’s *t*-test).

Next, we reviewed the putative secondary structure of *pspC* and *lph* RNA thermosensors using Vienna *RNAfold* package and identified secondary structures with occluded RBS (S7E and S7D Figs). The whole intergenic regions upstream of *pspC* and *lph* together with their respective first 90 coding bps were cloned into GFP-expressing plasmids pEGFP-N2 and transformed into *E. coli*. To eliminate any bacterial host factor involvement, these GFP-fusion constructs were subjected to *in vitro* transcription/translation at different biological relevant temperatures (S8 Fig). The PspC and PH thermosensors displayed gradual increment of expression, similar to the rheostat expression of their capsular counterparts CpsA and BcsA.

The finding of an elevated factor H binding proteins at higher temperatures from protein lysates are not definite evidence that more surface exposed proteins are produced, nor that FH binding occurs to the bacterial surface. To establish whether temperature could affect binding of FH, *S. pneumoniae* and *H. influenzae* grown at different temperatures were exposed to fluorescently labeled FH and analyzed by flow cytometry. *S. pneumoniae* showed an increased ability to bind FH when grown at 40°C, compared to 32°C (Fig 2C). Fluorescence microscopy staining was consistent in showing higher intensity of FH binding in *S. pneumoniae* grown at 40°C compared to 32°C (Fig 2E). Similarly, *H. influenzae* grown at different temperatures also showed differences in the flow cytometry analysis (Fig 2D). Two populations were observed when grown at 40°C as indicated by the two maxima in the histogram. When exposed to labeled FH, bacteria from this culture showed a clear increase in fluorescence intensity. Fluorescence microscopy staining revealed almost no fluorescence on bacteria grown at 32°C. When grown at 40°C, a subpopulation was found that was strongly positive for FH, while remaining bacteria were negative (Fig 2F). The microscopy observation is consistent with the two maxima in flow cytometry.

To test if higher expression of polysaccharide capsules and FH binding proteins at higher temperature could increase bacterial survival when facing human immune factors, serum killing and opsonophagocytosis assays were performed. *H. influenzae* was grown overnight at 30°C and pre-incubated at 30°C or 37°C for one hour prior to serum killing assays. *H. influenzae* pre-incubated at 37°C showed a significantly higher serum survival rate after 30 minutes compared to those at 30°C (Fig 3A). Pneumococci are naturally resistant to serum killing, therefore the protective role at different temperatures was studied using phagocytosis assays. *S. pneumoniae* was grown at either 30°C or 37°C, and opsonized with human serum prior to incubation with human THP-1 macrophages. Serum opsonized *S. pneumoniae* grown at 37°C was phagocytosed significantly less than those grown at 30°C (Fig 3B). Altogether the results show that at higher temperature, both *S. pneumoniae* and *H. influenzae* are better at evading complement mediated killing.

**Figure 3.**
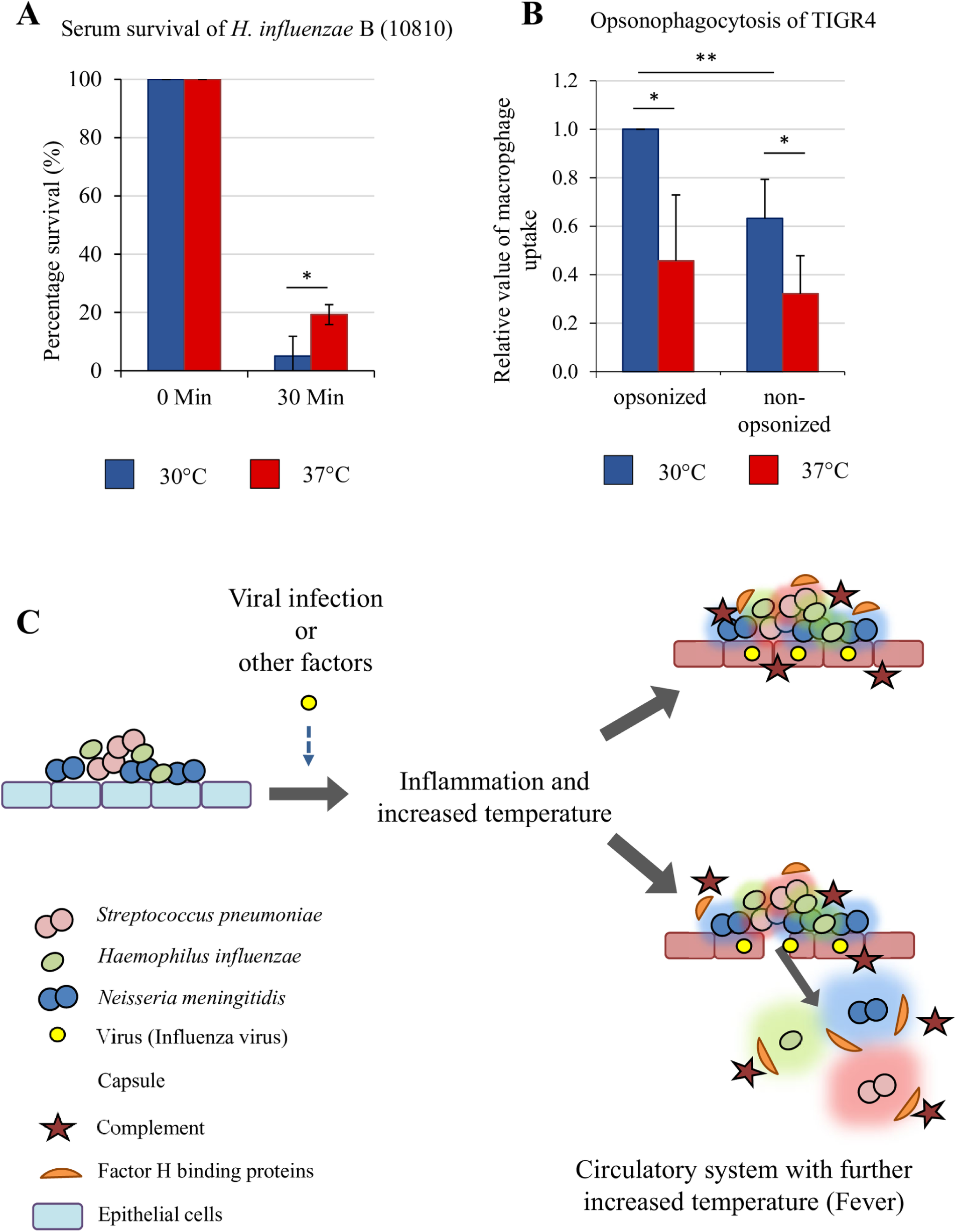
Temperature changes influence complement escape by *S. pneumoniae* and *H. influenzae*. A) *H. influenzae* were pre-incubated at 30°C or 37°C in RMPI for 1 hour prior to addition of human serum. The mixtures were then incubated at 37°C for 30 minutes. *H. influenzae* pre-incubated at 37°C shows more resistance to complement-mediated killing than those at 30°C. B) Opsonophagocytosis assay: *S. pneumoniae* TIGR4 was grown at 30°C or 37°C. The bacteria were then opsonized with human serum for 30min at 37°C. Non-opsonized bacteria were exposed to only RPMI for 30min at 37°C. Pneumococci grown at 37°C are more resistant to phagocytosis by macrophages than those grown at 30°C. Experiments were performed on three separate occasions and error bars denote s.e.m. * *p* < 0.05, ** p < 0.01 (Student’s *t*-test). C) Model of pathogenesis, from commensalism to systemic infection. Infection or other factors induce mucosal surface inflammation and a raise in temperature, leading to increased expression of virulence determinants such as capsule and FH binding proteins of *S. pneumoniae, H. influenzae* and *N. meningitidis*. This thermoregulation helps the bacteria to evade immune responses and enables better bacterial survival. The compromised integrity of the mucosal surfaces during viral replication serves as site of dissemination for bacteria into the circulatory system. Consequently, the temperature adapted bacteria will have better chance of surviving within warmer body sites, increasing the rate of systemic infections.

## Discussion

*S. pneumoniae* and *H. influenzae* are major causes of bacterial meningitis in children and adults[24]. These pathogens have evolved to colonize the human nasopharynx, a region replete with other bacteria and environmental stress factors. In addition, acute inflammation and pyretic response caused by systemic diseases such as influenza viruses could alter the local temperature within the nasopharynx[25]. It has been demonstrated that individuals with Influenza A virus infection are particularly susceptible to a secondary pneumococcal infection (reviewed by Brundage, J.F.)[26], but whether the fever response to the viral infection is a predisposing factor is currently unknown.

Elevation of temperature has been shown to act as a danger signal for *N. meningitidis*, prompting the bacterium to enhance immune evasion[14, 27]. Biosynthesis of the meningococcal sialic acid containing capsules and fHbp is governed by two independent RNA thermosensors. In agreement with this finding, we here show that *S. pneumoniae* and *H. influenzae* also are able to evade complement mediated killing by sensing temperatures via RNA molecules.

Through molecular and biochemical analyses of *S. pneumoniae* and *H. influenzae*, we identified four sequence unique RNA thermosensors controlling the expression of their capsular gene (Cps and Bcs) and FH binding proteins (PspC and PH). Previous studies have shown that mutations within the 5′-region of *cps* could disrupt the production of the capsule[23, 28]. However, these authors did not fully understand mechanisms leading to disruption of the capsule production. Our findings here explain that the mutations generated would stabilize the RNA thermosensor’s secondary structure thus resulting in lower capsule production. Like all other characterized RNA thermosensors[14, 20], we were able to show that single nucleotide mutations in the *cps* 5′-UTR could disrupt its thermosensing abilities. Furthermore, we found that the nucleotide sequences within the 5′-UTR of *cps*, essential for forming the RNA secondary structure, are highly conserved among various pneumococcal strains of different capsular serotypes. Naturally occurring polymorphisms in the *cps* 5′-UTR of serotypes 1, 2 and 22F could neither alter the putative RNA secondary structures, nor the thermoregulation ability of Cps (Fig S6).

Our results show that the increased translation of pneumococcal CpsA at higher temperature indeed led to higher production of capsule. Studies have previously shown that capsular shedding enable pneumococci to evade complement killing and facilitate bacterial spread[29, 30]. These shed capsules are immunogenic and free floating components that could additionally sequester antibodies and immune cells. In agreement with these studies, we also observed more capsule in the supernatant from pneumococci grown at higher temperature.

Temperature mediated production of capsules and binding of human FH in *S. pneumoniae* and *H. influenzae* could indicate better protection for the bacteria from immune killing. Our work here demonstrates that both bacteria are indeed able to evade complement mediated killing at higher temperature as shown using assays for serum killing and opsonophagocytosis. An increase in temperature triggered by inflammation in the host may act as a ‘danger signal’ for *S. pneumoniae* and *H. influenzae* priming their defenses against the recruitment of immune effectors onto the mucosal surface. While attuned to higher threat, it remains largely unknown how these pathogens breach from the mucosa into the bloodstream and further into the brain. We hypothesize that the nasopharyngeal tissue could be damaged during prior Influenza A viral infections, serving as an entry site for the bacteria. Temperature increment caused by a local inflammation by primed pathogens could enhance evasion of immune responses and increase bacterial growth, leading to bacteremia and the pathogen crossing the blood brain barrier to cause meningitis[31-33]. A proposed infection model influenced by temperature is shown in Fig 3C.

The expression of capsular polysaccharides and FH binding proteins is an important survival strategy for these meningitis causing pathogens. However, overexpression of such factors at certain sites, such as on nasopharyngeal mucosal surfaces, could evoke undesired immune reactions for the bacteria, thus jeopardizing colonization. It has previously been demonstrated that high capsular expression in pneumococci may have negative effects on adhesion to the respiratory mucosa[34]. Thermosensor mediated modulation of capsular expression may therefore be a central strategy for these meningitis pathogens to optimally colonize their normal habitat, the human nasopharynx. At the same time, and when necessary (i.e. local inflammation), these bacteria could rapidly express certain properties in order to avoid immune killing.

We here reveal that human restricted nasopharyngeal pathogens, *S. pneumoniae* and *H. influenzae*, sense ambient temperature changes. RNA thermosensors positively influence the expression of their major virulence determinants at warmer growth conditions. Interestingly, the four novel RNA thermosensors described here, together with the two known neisserial RNA thermosensors, do not possess any sequence similarity among them, but all retain the same thermosensing ability. This suggests that while nucleotide sequences within the 5′-UTR could be dispensable, a functional RNA thermosensor imperatively depends on its secondary structure with a weakly base-paired RBS. It is most likely that these RNA thermosensors in *S. pneumoniae, H. influenzae* and *N. meningitidis* have evolved independently to sense the same temperature cue in the nasopharyngeal niche to avoid immune killing.

RNA-mediated virulence gene regulation is still an understudied phenomenon. However, with recent RNA sequencing methods such as RNA structurome sequencing analysis[35], comprehensive RNA thermosensor maps could be generated. In addition, nuclear magnetic resonance (NMR) spectroscopy could be used to investigate the RNA structures[36-38]. We believe that such studies will provide valuable information on the functional importance and dynamics on RNA regulation in these pathogens, ultimately contributing to the understanding of bacterial pathogenesis.

## Materials and methods

### Bacterial strains, plasmids and growth conditions

Strains, oligonucleotides and plasmids are listed in Table 1. *Streptococcus pneumoniae* were grown on blood agar plates, *Haemophilus influenzae* grown on chocolate agar plates and *Neisseria meningitidis* grown on BHI agar (1% w/v) supplemented with Levinthal’s base at 37°C and 5% CO_2_ overnight. *E. coli* was grown in Luria–Bertani (LB) agar at 37 °C overnight When necessary, antibiotic was added to the following final concentrations: kanamycin, 50 µg ml^-1^. In order to generate the CpsA-GFP, BscA-GFP, CssA-GFP, PspC-GFP and LpH-GFP constructs in the plasmid pEGFP-N2[14], the following oligonucleotides were used to amply the fragments from their respective gDNAs; CspA(GFP)-F with CpsA(8C-GFP)-R, BscA(GFP)-F with BscA(8C-GFP)-R, CssA(GFP)-F with CssA(8C-GFP)-R, PspC(GFP)-F with PspC(30C-GFP)-R and LpH(GFP)-F with LpH(30C-GFP)-R. PCR fragments together with pEGFP-N2 plasmid were then digested with *EcoRI* and XmaI restriction enzymes (New England Biolab) according to manufacturer’s protocol and ligated using T4 ligase (Promega). The resulting constructs were sequenced to ensure that no other changes had occurred prior transformation into XL10-Gold *E.coli* (Agilent).

**Table 1.**
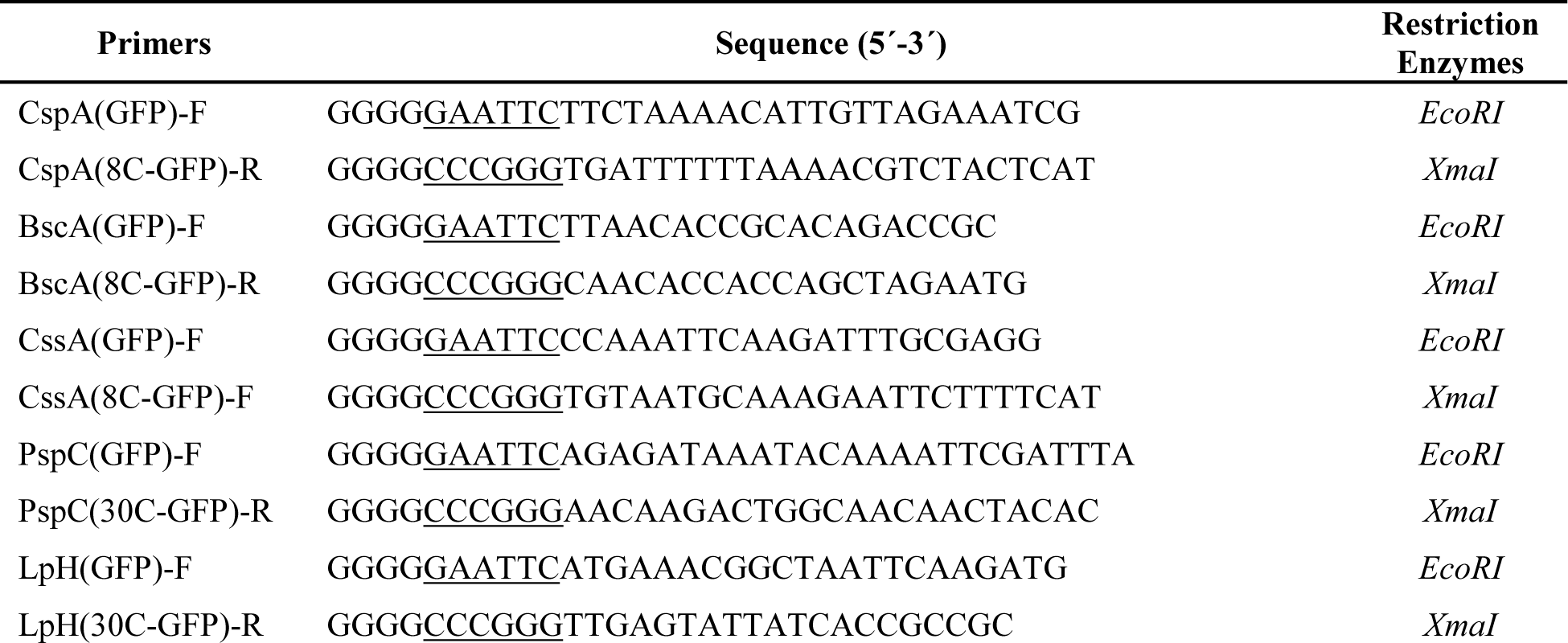
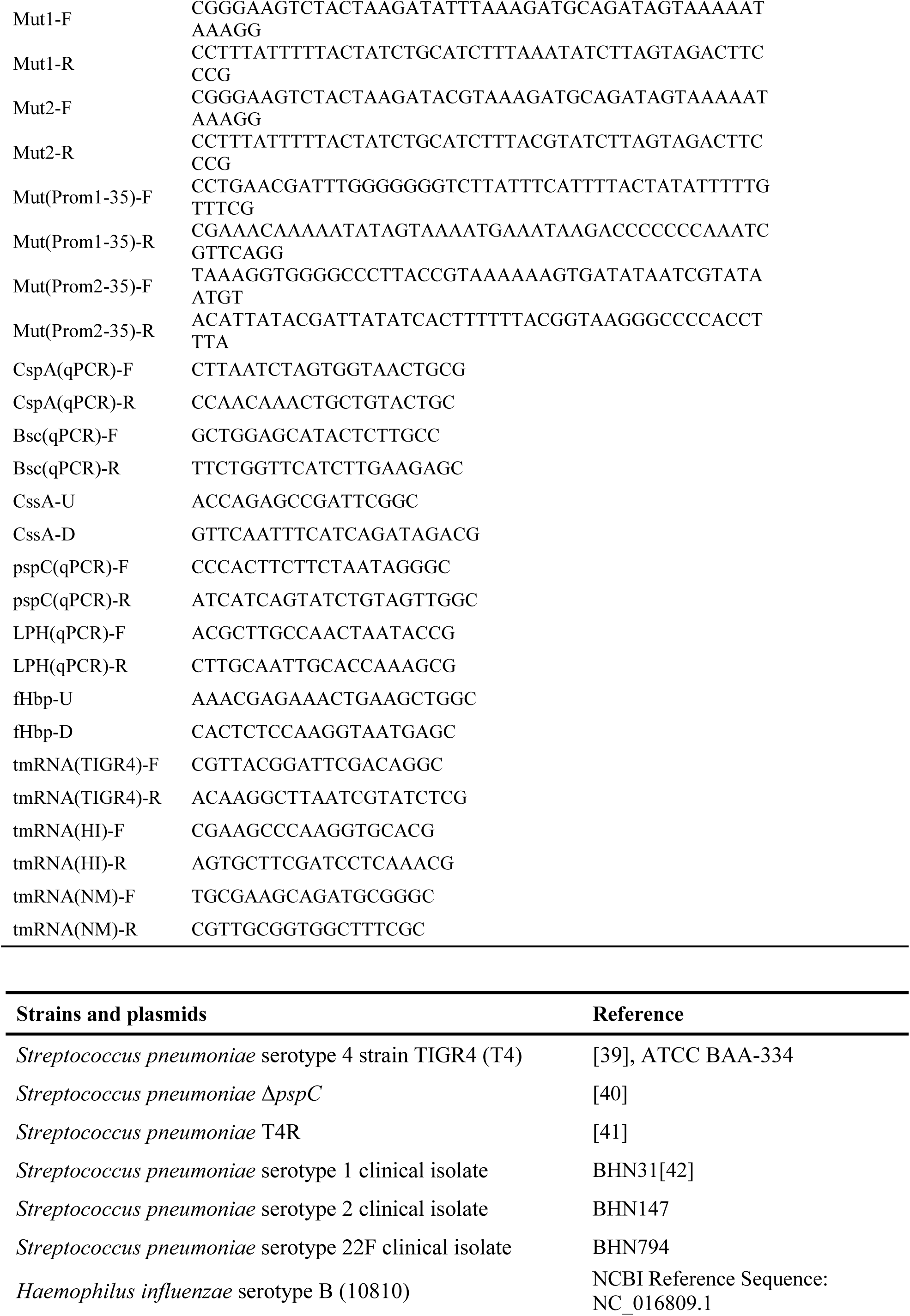
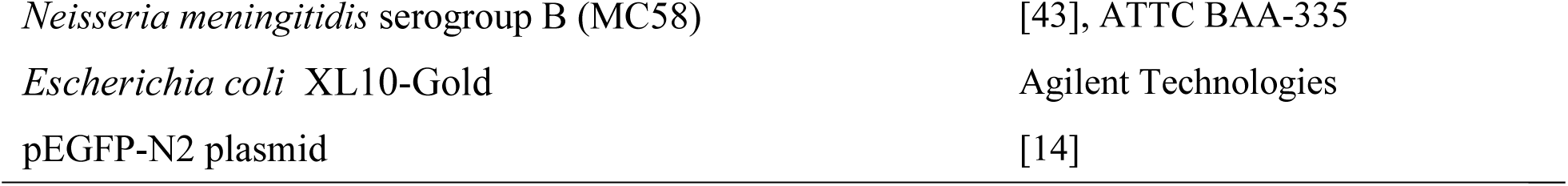
Oligonucleotides, strains and plasmids

### Site-Directed Mutagenesis

In order to obtain site-specific mutations in CpsA-GFP constructs, we used the Phusion High-Fidelity DNA Polymerases (Thermo Scientific) for site-directed mutagenesis. A plasmid harboring the CpsA-GFP fragment was used as template in the mutagenesis reactions. To achieve disrupted thermosensing, CpsA mut-1 and mut-2, oligonucleotide pairs of Mut1-F with Mut1-R and Mut2-F with Mut2-R respectively, were used (Table 1). To achieve promoter-1 and promoter-2 mutants of CpsA-GFP, oligonucleotide pairs of Mut(Prom1-35)-F with Mut(Prom1-35)-R and Mut(Prom2-35)-F with Mut(Prom2-35)-R respectively, were used. The resulting mutant constructs were sequenced to ensure that no other changes had occurred.

### RNA isolation and Northern blotting

*S. pneumoniae, H. influenzae* and *N. meningitidis* were grown in liquid culture to an OD_600 nm_ of ∼0.5 (5×10^8^ CFUs/mL; 2.5×10^9^ CFUs/mL; and 2×10^9^ CFUs/mL respectively) before RNA extraction. RNA was isolated using the FastRNA^®^ Pro Blue Kit (MP Biomedical) following the manufacturer’s protocol. Northern blot analyses were performed according to standard protocols. Briefly, PCR probes were generated using oligonucleotide pairs CpsA(qPCR)-F with CpsA(qPCR)-R, Bsc(qPCR)-F with Bsc(qPCR)-R, CssA-U with CssA-D, pspC(qPCR)-F with pspC(qPCR)-R, LPH(qPCR)-F with LPH(qPCR)-R, fhbp-U with fhbp-D, tmRNA(TIGR4)-F with tmRNA(TIGR4)-R, tmRNA(HI)-F with tmRNA(HI)-R and tmRNA(NM)-F with tmRNA(NM)-R. PCR fragments were then labelled using Megaprime DNA Labeling Systems (GE Healthcare). Primers used are listed in Table 1.

### *In vitro* transcription/translation

Plasmids containing the desired constructs were *in vitro* transcribed in an *E. coli* S30 Extract system for Linear Templates kit (Promega) according to the manufacturer’s instructions. In brief, *cpsA-gfp, bscA-gfp, cssA-gfp, pspC-gfp, lph-gfp* and *prfA-gfp* plasmids (for plasmid constructs see bacterial strains, plasmids and growth conditions) were digested using NotI restriction enzyme and purified using QiAquick PCR purification kit (Qiagen). One microgram of *cpsA-gfp, bscA-gfp, cssA-gfp, pspC-gfp, lph-gfp* and *prfA-gfp* digested plasmids mixtures were incubated at 30°C, 32°C, 34°C, 36°C, 38°C and 40°C for 1h before transferring onto ice for 5 min. Samples were acetone-precipitated, re-suspended in 1X sample buffer, and separated on a 12% polyacrylamide gel before being transferred onto Trans-Blot® Turbo™ Midi (PVDF) membranes (Biorad) using the Trans-Blot^®^ Turbo™ Transfer System (Biorad). Membranes were developed following the protocol of the ECL western blotting kit (Amersham), using anti-GFP (BD-living colours) as primary antibody and an HRP-conjugated anti-mouse as the secondary antibody (GE Healthcare).

### SDS–PAGE and Western blotting

*S. pneumoniae, H. influenzae* and *N. meningitidis* were grown in liquid culture to an OD_600 nm_ of ∼0.5 before protein isolation. Total protein was isolated from whole cell lysates using BugBuster® (Merck). Total protein concentrations were quantified by Pierce BCA quantification kit (Thermo Fisher Scientific). 20µg of protein was loaded into each lane of the gel. Western blot analyses were performed according to standard protocols. For western blot analysis, membranes were washed three times in 0.05% (w/v) dry milk/PBS with 0.05% (v/v) Tween-20 for 10 min, and then incubated with the primary antibody for 1 h. Membranes were washed again three times and incubated for a further hour with a secondary, horseradish peroxidase (HRP)-conjugated antibody for 1 h. Binding was detected with an ECL Western Blotting Detection kit (Amersham) and exposed to ECL Hyperfilm. Anti-CpsB rabbit antibody (gift from Professor James C. Paton) was used at a dilution of 1:1000. Anti-CbpA/PspC rabbit antibody (gift from Professor Elaine I. Tumanen) was used at a dilution of 1:8000. Anti-Lph rabbit antibody (gift from Professor Kristian Reisbeck) was used at a dilution of 1:5000. Anti-fHbp Anti-RecA rabbit antibody (Abcam) was used at a dilution of 1:8000. Anti-LytA rabbit antibody was used at a dilution of 1:2000. Anti-GAPDH rabbit antibody was used at a dilution of 1:2000. Anti-fHbp mouse antibody was used at a dilution of 1:5000. Anti-GFP mouse antibody (BD living colors) was used at a dilution of 1:8000. Anti-Kanamycin rabbit antibody (Novus) was used at a dilution of 1:1000. For Far western, Complement factor H from human plasma (Sigma Aldrich) was used at a dilution of 1:10000 prior to Anti-Factor H goat antibody at a dilution of 1:1000. HRP-conjugated anti-mouse, anti-rabbit or anti-goat as the secondary antibody (GE Healthcare) were used at a dilution of 1:3000.

### Serum killing assay

To determine the effect of temperature on complement sensitivity, *H. influenzae* was grown in liquid media according to our previous published protocol^13^. In brief, *H. influenzae* strain was grown on chocolate agar plates overnight at 30°C and re-suspended in PBS. Bacteria were diluted to a final concentration of 1 × 10^7^ c.f.u. ml^-1^ in RPMI medium (Gibco). To compare the sensitivity of bacteria at different temperatures, resuspended bacteria in RPMI were then split and incubated at 30°C and 37°C for a further 1 h. One-million c.f.u. were incubated with 25% human immune serum (Sigma-Aldrich) for 30 min, and the proportion of bacteria surviving was determined by plating 10 μl aliquots onto chocolate plates and counting the number of colonies after overnight incubation; differences were analysed with the Student’s *t*-test.

### Opsonophagocytosis assay

*S. pneumoniae* was grown on blood agar plates at 30°C or 37°C and 5% CO_2_ overnight. Colonies were inoculated into C+Y medium and grown until exponential phase (∼5×10^8^ CFUs/mL) at 30°C or 37°C prior to incubation with THP-1 macrophages. Serial dilutions were plated on blood agar plates and incubated over night at 37°C and 5% CO_2_ to determine the number of phagocytosed bacteria.

### Polysaccharide dot blot

*S. pneumoniae* was grown in C + Y media from −80°C 20% glycerol stocks, prepared from cultures in mid-logarithmic phase. Bacteria were grown at 30°C, 32°C, 37°C, or 40°C until cultures reached 5×10^8^ CFUs/mL. Cells and medium were split by centrifugation and supernatant isolated. 5µL pure supernatant were blotted on PVDF membrane (Hybond-P, Amersham Corp.) and dried for 1h at 37°C. Purified type 4 capsular polysaccharide (Statens Serum Institute, Denmark) was used as positive control. Capsule components of *S. pneumoniae* TIGR4 were detected using 1:2 optimized rabbit type-4 antiserum (Statens Serum Institute) and 1:20,000 anti-Rabbit-HRP. Optimized antiserum was produced by exposing live T4R (non-encapsulated TIGR4) to Type 4 antiserum, 1:100 diluted in PBS for 30 min at 37°C.

### Flow cytometry analysis and fluorescence microscopy

Bacteria were grown in liquid from −80°C 20% glycerol stocks, prepared from cultures in mid-logarithmic phase. *S. pneumoniae* grew in C + Y media, and H. *influenzae* in BHI media at 30°C, 32°C, 37°C, or 40°C until cultures reached 5×10^8^ (*S. pneumoniae*) or 2.5×10^9^ (*H. influenzae*) CFU/mL. Capsule of *S. pneumoniae* TIGR4 was detected using 1:2 optimized rabbit type-4 antiserum and 1:100 anti-Rabbit-Alexa-Fluor-488 (Thermo Fisher). Binding of human FH (Merck Millipore) was detected with Alexa-Fluor-488 labeled human FH. Labeling was performed with the Alexa Fluor™ 488 Microscale Protein Labeling Kit. Sample data was acquired on Gallios flow cytometer (Beckman Coulter) and at least 2×10^5^ events recorded. Fluorescence intensity was analyzed using Kaluza software (Beckman Coulter). For fluorescence microscopy 5µL of the samples were dried on cover slip, then covered with Vectashield H-1400 and analyzed using the DeltaVision Elite (GE Healthcare).

## Acknowledgements

This work was supported by grants from the Knut and Alice Wallenberg Foundation, the Swedish Foundation for Strategic Research (SSF), the Swedish Research Council (to EL and BHN), ALF grant from Stockholm County Council, and Karolinska Institutet. We are grateful to J. Johansson and S. Normark for scientific discussions.

## Author contribution

H.E., L.S. and E.L. performed the experiments. H.E., L.S., A.P., B.H-N. and E.L. analysed the data. B.H-N. and E.L. provided overall directions. H.E. performed literature search and wrote the manuscript with input from B.H-N. and E.L.

## Supporting information

**S1 Fig.**

A) Promoters and 5′-UTR sequences of the capsular biosynthesis of *cpsA* of TIGR4 S. *pneumoniae*. Native promoter (red), thermosensing promoter (green).

B) Promoter and 5′-UTR sequences of the capsular biosynthesis of *H. influenzae* (*bcsA*). Promoter (green)

(A and B): transcriptional start site (arrows), ribosome binding site (RBS) (blue), coding region (shaded box).

C) Thermoregulation of CssA- and PrfA-GFP fusion protein is detected in *E. coli* by western blot analysis. RecA and Neomycin-phosphotransferase (Npt) are used as loading controls.

**S2 Fig.**

A) Promoters and 5′-UTR sequences of the capsular biosynthesis of *cpsA* of TIGR4 S. *pneumoniae*. Native promoter/promoter-2 (grey box), thermosensing promoter/promoter-1 (blue box), predicted transcriptional start site (arrows), ribosome binding side (RBS) (blue), coding region (red box). Purple sequences denote mutations within −35 element of each promoter.

B) Western blot analysis shows that thermoregulation of CpsA is disrupted in a Promoter-1 mutant (Prom-1 Mut) whereas a mutant in Promoter-2 (Prom-2 Mut) retains its thermosensing ability. Site-directed mutageneses were performed to replace the −35 sequences of each promoter. In Prom-1 Mut background, only promoter 2 is active and in the Prom-2 Mut background, only promoter 1 is active. RecA was used as a loading control.

**S3 Fig.**

Northern blot analyses demonstrate that *cpsA, pspC, bcsA* and *lph* mRNA transcripts are not thermoregulated. (*tmRNA*, transfer-messenger RNA, *N. meningitidis cssA* and *fHbp* used as controls).

**S4 Fig.**

*In vitro* transcription/translation assay comparing capsule producing enzymes with known thermosensors at a range of biological temperatures. CpsA and BcsA show thermosensing. *N. meningitidis* CssA and *L. monocytogenes* PrfA thermosensor are used as positive controls. Blots are representative of experiments performed on at least three occasions.

**S5 Fig.**

A) Predicted RNA secondary structure of the whole 5′-UTR and 24 bases of *cpsA*-mRNA.

B) Predicted RNA secondary structure of the whole 5′-UTR and 24 bases of *bcsA*-mRNA.

(A and B): Ribosome binding site (RBS) indicated in blue, start codon in green, putative structures predicted with *RNAfold* program and visualized with *VARNA* applet. Length of 5′-UTRs in nucleotides (nts) and ΔG values as indicated.

**S6 Fig.**

A) Thermoregulation of pneumococcal capsular protein B (CpsB) and FH binding protein (PspC) is evident in *S. pneumoniae* across different serotypes and lineages. CC, clonal complex; ST, sequence type.

B) 5′-UTR sequence alignment of Serotype 1, 2, 4 and 22F using *CLUSTAL O(1.2.4) multiple sequence alignment*;

C-E) Predicted RNA secondary structures of the whole 5′-UTR and 24 bases of *cpsA*-mRNA. Serotype 1 (C), serotype 2 (D) and serotype 22F (E).

(**A** to **E**): ribosome binding site (RBS) indicated in blue, start codon in green, mutations compared to TIGR4 in red, secondary structures predicted by *RNAfold* and visualized with *VARNA* applet.

**S7 Fig.**

A) Promoters and 5′-UTR sequences of *pspC* in *S. pneumoniae.*

B) Promoters and 5′-UTR sequences of *lph* in *H. influenzae*

(A and B): promoter (green), transcriptional start site (arrows), ribosome binding site (RBS) (blue), coding region (shaded box).

C) Predicted RNA secondary structure of the 5′-UTR and 90 bases of *pspC*-mRNA.

D) Predicted RNA secondary structure of the 5′-UTR and 90 bases of *lph*-mRNA.

(C and D): RBS indicated in blue, start codon in green, start codon in green, putative secondary structure predicted with *RNAfold* and visualized with *VARNA* applet. Length of 5′-UTRs in nucleotides (nts) and ΔG values as indicated.

**S8 Fig.**

*In vitro* transcription/translation assay comparing capsule producing enzymes with known thermosensors at a range of biological temperatures. PspC and PH show thermosensing. *N. meningitidis fHbp* and *L. monocytogenes* PrfA thermosensors are used as positive controls. Blots are representative of experiments performed on at least three occasions.

