## Supplementary Figures for "Meningitis pathogens evade immune responses by thermosensing"

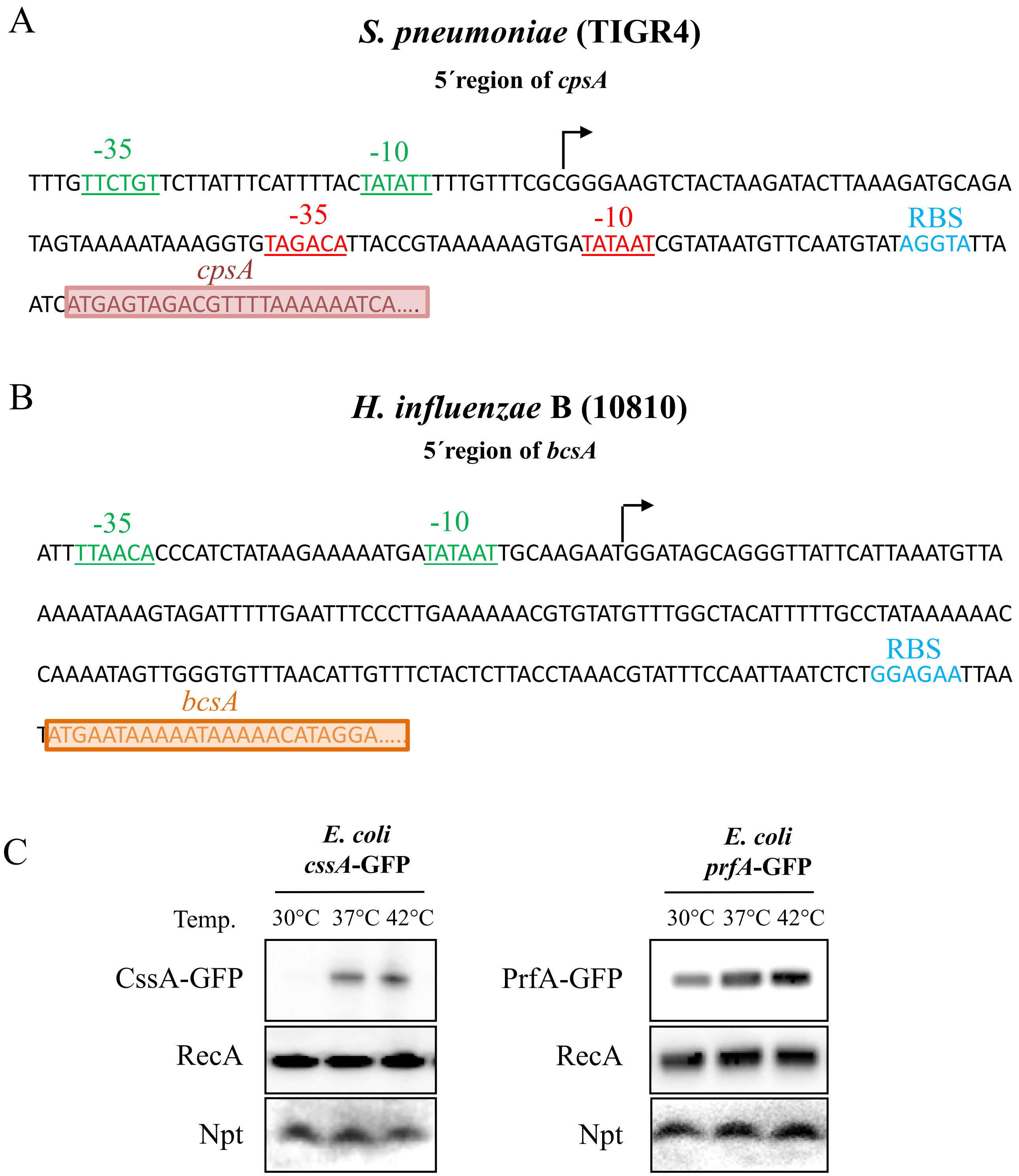
A
S. pneumoniae (TIGR4)
5´region of cpsA
-35
-10
TTTGTTCTGTTCTTATTTCATTTTACTATATTTTTGTTTCGCGGGAAGTCTACTAAGATACTTAAAGATGCAGA
-35
-10
RBS
TAGTAAAAATAAAGGTGTAGACATTACCGTAAAAAAGTGATATAATCGTATAATGTTCAATGTATAGGTATTA
cpsA
ATCATGAGTAGACGTTTTAAAAAATCA….
B
H. influenzae B (10810)
5´region of bcsA
-35
-10
ATTTTAACACCCATCTATAAGAAAAATGATATAATTGCAAGAATGGATAGCAGGGTTATTCATTAAATGTTA
AAAATAAAGTAGATTTTTGAATTTCCCTTGAAAAAACGTGTATGTTTGGCTACATTTTTGCCTATAAAAAAC
RBS
CAAAATAGTTGGGTGTTTAACATTGTTTCTACTCTTACCTAAACGTATTTCCAATTAATCTCTGGAGAATTAA
bcsA
TATGAATAAAAAATAAAAACATAGGA….
C
E. coli cssA-GFP
E. coli prfA-GFP
Temp.
30°C 37°C 42°C
30°C 37°C 42°C
CssA-GFP
PrfA-GFP
RecA
RecA
Npt
Npt

**S1 Fig.**

**S1 Fig.**

A) Promoters and 5´-UTR sequences of the capsular biosynthesis of *cpsA* of TIGR4 S. *pneumoniae*. Native promoter (red), thermosensing promoter (green).

B) Promoter and 5´-UTR sequences of the capsular biosynthesis of *H. influenzae* (*bcsA*). Promoter (green)

(A and B): transcriptional start site (arrows), ribosome binding site (RBS) (blue), coding region (shaded box).

C) Thermoregulation of CssA- and PrfA- GFP fusion protein is detected in *E. coli* by western blot analysis. RecA and Neomycin-phosphotransferase (Npt) are used as loading controls.

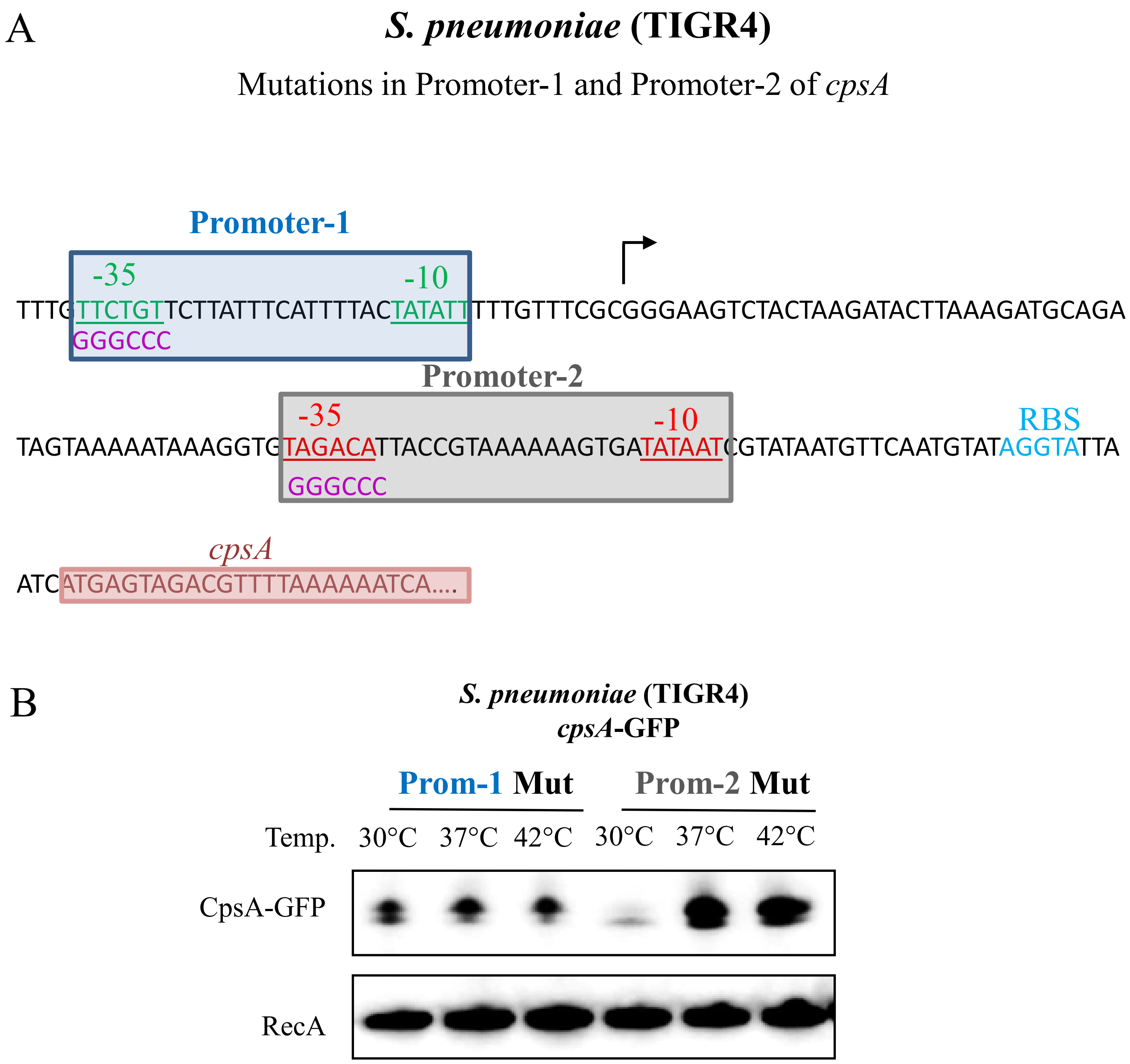

A
S. pneumoniae (TIGR4)
Mutations in Promoter-1 and Promoter-2 of cpsA
Promoter-1
-35
-10
TTTGTTCTGTTCTTATTTCATTTTACTATATTTTTGTTTCGCGGGAAGTCTACTAAGATACTTAAAGATGCAGA
GGGCCC
Promoter-2
-35
-10
RBS
TAGTAAAAATAAAGGTGTAGACATTACCGTAAAAAAGTGATATAATCGTATAATGTTCAATGTATAGGTATTA
GGGCCC
cpsA
ATCATGAGTAGACGTTTTAAAAAATCA....
B
S. pneumoniae (TIGR4)
cpsA-GFP
Prom-1 Mut
Prom-2 Mut
Temp. 30°C 37°C 42°C 30°C 37°C 42°C
CpsA-GFP
RecA


**S2 Fig.**

**S2 Fig.**

A) Promoters and 5´-UTR sequences of the capsular biosynthesis of *cpsA* of TIGR4 S. *pneumoniae*. Native promoter/promoter-2 (grey box), thermosensing promoter/promoter-1 (blue box), predicted transcriptional start site (arrows), ribosome binding side (RBS) (blue), coding region (red box). Purple sequences denote mutations within -35 element of each promoter.

B) Western blot analysis shows that thermoregulation of CpsA is disrupted in a Promoter-1 mutant (Prom-1 Mut) whereas a mutant in Promoter-2 (Prom-2 Mut) retains its thermosensing ability. Site-directed mutageneses were performed to replace the -35 sequences of each promoter. In Prom-1 Mut background, only promoter 2 is active and in the Prom-2 Mut background, only promoter 1 is active.  RecA was used as a loading control.

Northern Blots

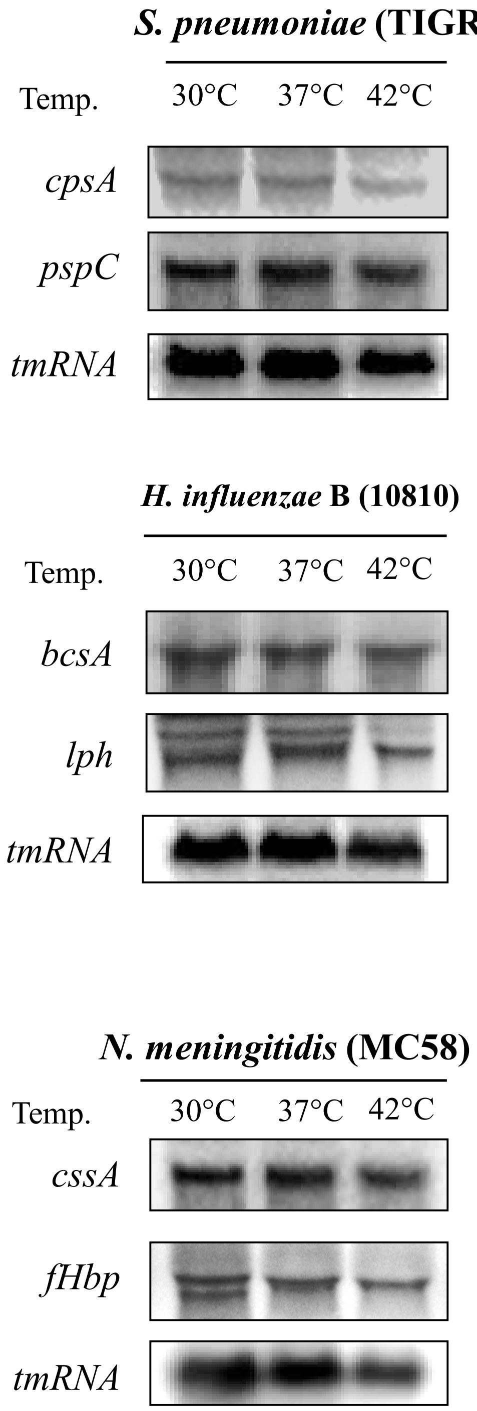


**S3 Fig.**

**S3 Fig.**

Northern blot analyses demonstrate that *cpsA*, *pspC*, *bcsA* and *lph* mRNA transcripts are not thermoregulated. (*tmRNA*, transfer-messenger RNA, *N. meningitidis cssA* and *fHbp* used as controls).

A

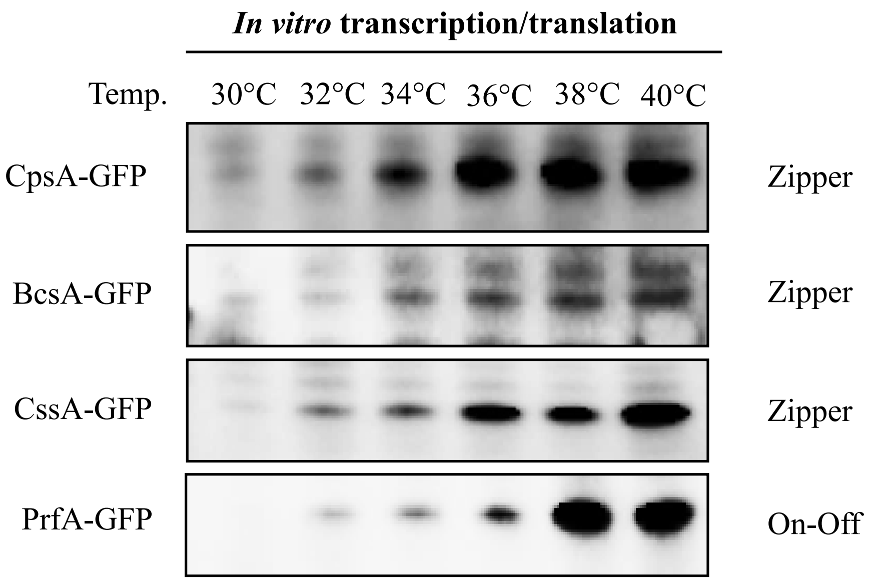
*In vitro* transcription/translation
Temp. 30°C 32°C 34°C 36°C 38°C 40°C
CpsA-GFP
Zipper
BcsA-GFP
Zipper
CssA-GFP
Zipper
PrfA-GFP
On-Off

**S4 Fig.**

**S4 Fig.**

*In vitro* transcription/translation assay comparing capsule producing enzymes with known thermosensors at a range of biological temperatures. CpsA and BcsA show thermosensing. *N. meningitidis* CssA and *L. monocytogenes* PrfA thermosensor are used as positive controls. Blots are representative of experiments performed on at least three occasions.

A

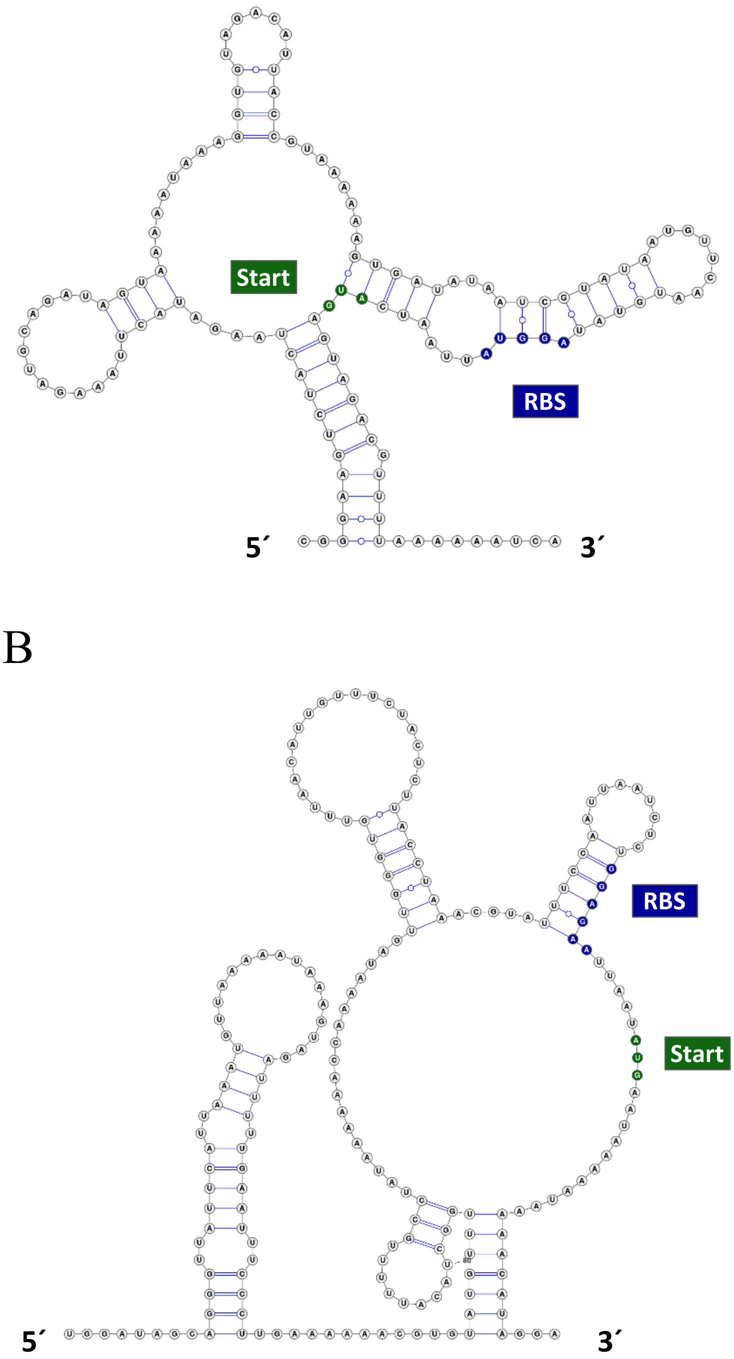


*cpsA* **5´-UTR**

133 nts

**ΔG = -19.08** kcal/mol

*bcsA* **5´-UTR**

198 nts

**ΔG = -29.70** kcal/mol

**S5 Fig.**

**S5 Fig.**

A) Predicted RNA secondary structure of the whole 5´-UTR and 24 bases of *cpsA*-mRNA.

B) Predicted RNA secondary structure of the whole 5´-UTR and 24 bases of *bcsA*-mRNA.

(A and B): Ribosome binding site (RBS) indicated in blue, start codon in green, putative structures predicted with *RNAfold* program and visualized with *VARNA* applet. Length of 5´-UTRs in nucleotides (nts) and ΔG values as indicated.

A

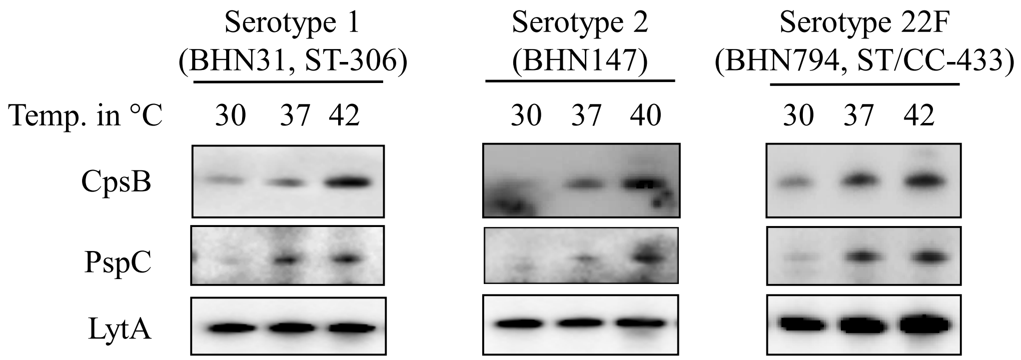


B

```
BHN147(Serotype-2)     CGGGAAGTCTACTAAGATACTTAAAGATGCAGATAGTGAA--AAAAGGTGTAGACATTAC  58
TIGR4(Serotype-4)      CGGGAAGTCTACTAAGATACTTAAAGATGCAGATAGTAAAAATAAAGGTGTAGACATTAC  60
BHN31(Serotype-1)      CGGGAAGTCTACTAAGATACTTAAAGATGCAGATAGTGAAAATAAAGGTGTAGACATTAC  60
BHN794(Serotype-22F)   CGGGAAGTCTACTAAGATACTTAAAGATGCAGATAGTGAAAATAAAGGTGTAGACATTAC  60
                       ************************************* **   *****************

BHN147(Serotype-2)     CGTAAAAAAGTGATATAATCGTAAGATGTTCAATGTATAGGTGTTAATCATGAGTAGACG  118
TIGR4(Serotype-4)      CGTAAAAAAGTGATATAATCGTATAATGTTCAATGTATAGGTATTAATCATGAGTAGACG  120
BHN31(Serotype-1)      CGTAAAAAAGTGATATAATTGTATGATGTTCAAGGTATAGGTGTTAATCATGAGTAGACG  120
BHN794(Serotype-22F)   CGTAAAAAAGTGATATAATCGTATGATGTTCAATGTATAGGTGTTAATCATGAGTAGACG  120
                       ******************* ***  ******** ******** *****************
                                                             RBS          Start

BHN147(Serotype-2)     TTTTAAAAAATCA 131
TIGR4(Serotype-4)      TTTTAAAAAATCA 133
BHN31(Serotype-1)      TTTTAAAAAATCA 133
BHN794(Serotype-22F)   TTTTAAAAAATCA 133
                       *************
```

**S6 Fig.**

C

**BHN31**

***cpsA* 5´-UTR**

133 nts

**ΔG = -17.50** kcal/mol

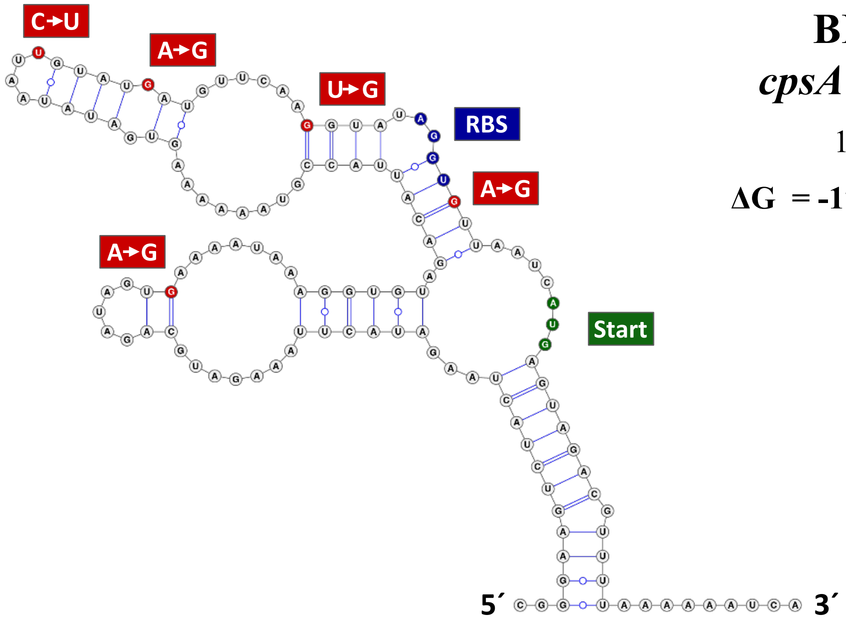


D

**BHN147**

***cpsA* 5´-UTR**

131 nts

**ΔG = -20.21** kcal/mol

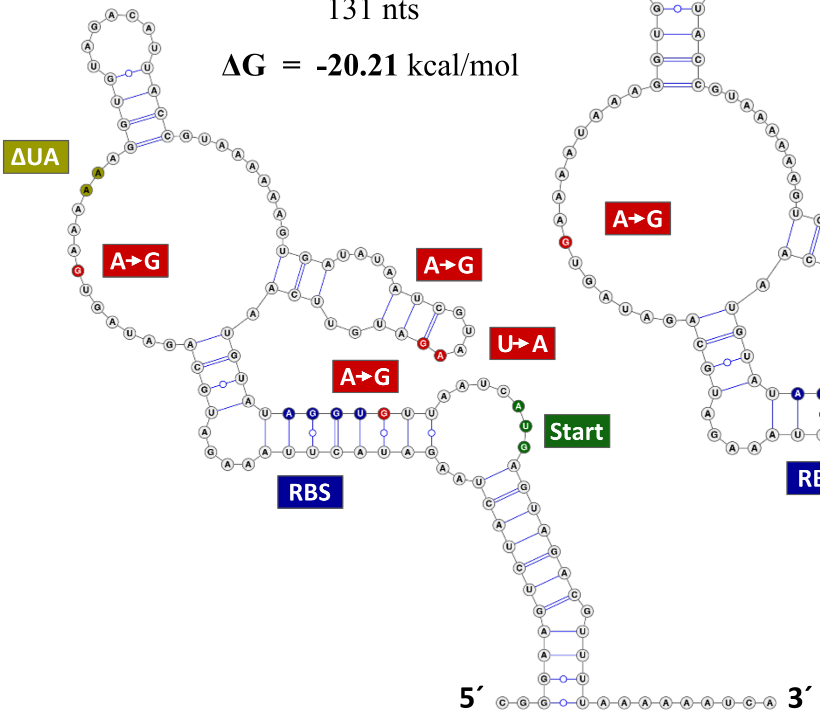


E

**BHN794**

***cpsA* 5´-UTR**

133 nts

**ΔG = -19.93** kcal/mol

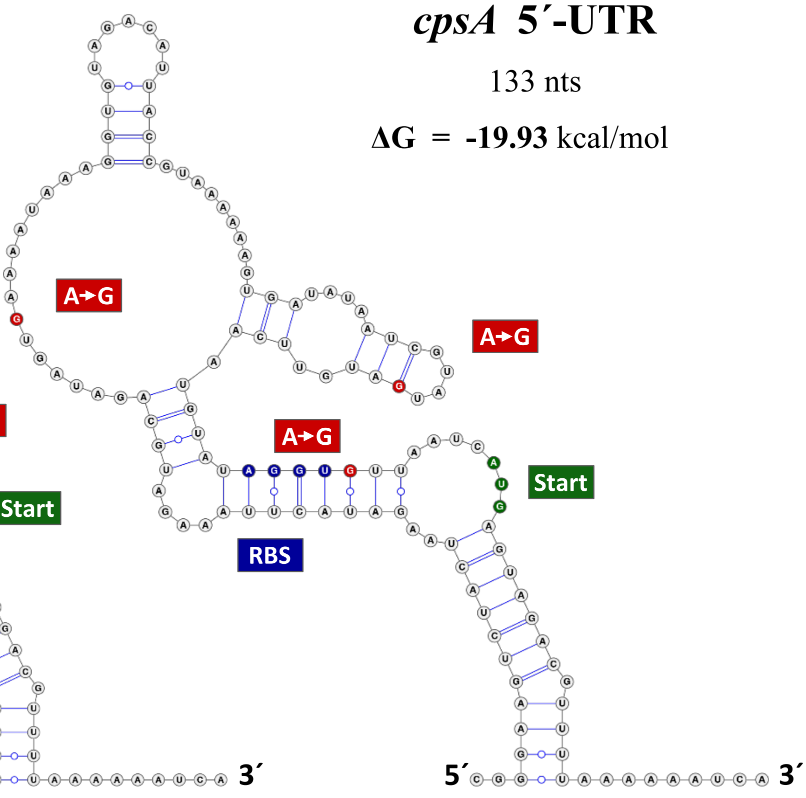


**S6 Fig.**

**S6 Fig.**

A) Thermoregulation of pneumococcal capsular protein B (CpsB) and FH binding protein (PspC) is evident in *S. pneumoniae* across different serotypes and lineages. CC, clonal complex; ST, sequence type.

B) 5´-UTR sequence alignment of Serotype 1, 2, 4 and 22F using *CLUSTAL O(1.2.4) multiple sequence alignment*;

C-E) Predicted RNA secondary structures of the whole 5´-UTR and 24 bases of *cpsA*-mRNA. Serotype 1 (C), serotype 2 (D) and serotype 22F (E).

(**A** to **E**): ribosome binding site (RBS) indicated in blue, start codon in green, mutations compared to TIGR4 in red, secondary structures predicted by *RNAfold* and visualized with *VARNA* applet.

A

*S. pneumoniae,* TIGR4 (*pspC*- SP_2190)

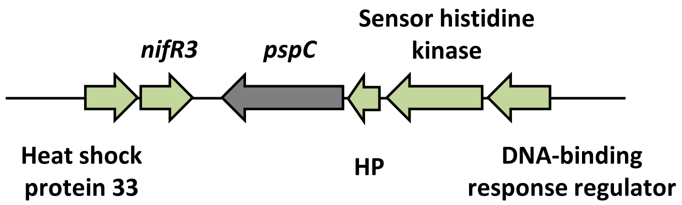


-35 TTTATA; -10 TGATAT

AGAGATAAATACAAAATTCGATTTATATACAGTTCATATTGAAGTGATATA

RBS AGGAA

GTAAGGTTAAAGAAAAAATATAGAAGGAAATAAACATGTTTGCATCAA

*pspC*

AAAGCGAAAGAAAAGTACATTATTCAATTCGTAAATTTAGTGTTGGAGT

AGCTAGTGTAGTTGTTGCCAGTCTTGTT.....

B

*H. influenzae* B, 10810 (*lph*- HIB_11110)

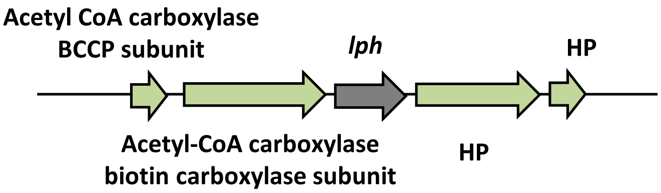


-35 TTTGAA

TTCATTTTATCCCAATGATTTATAAAATGTATTTGAAATGATATAGAATCC

-10 TATAAT; RBS AAGAT

TTATATAATTTGTAACTATTTTTTTTACTTAAGATAAAAAAACAATGAAA

*lph*

CTTAATCTTTCGAAATTCAGCTTAACTATTCTTACAACGGTAATGCTGG

CTTCCTGTGGAAGTGGCGGCGGTGATAATACTCAA.....

**S7 Fig.**

C

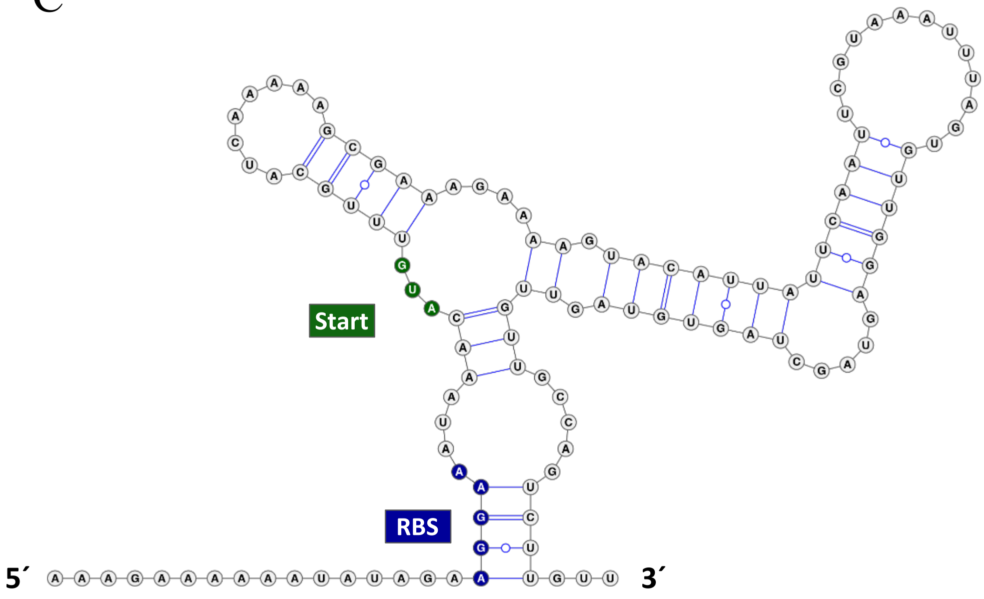


*pspC* **5´-UTR**

117 nts

**ΔG = -14.93** kcal/mol

D

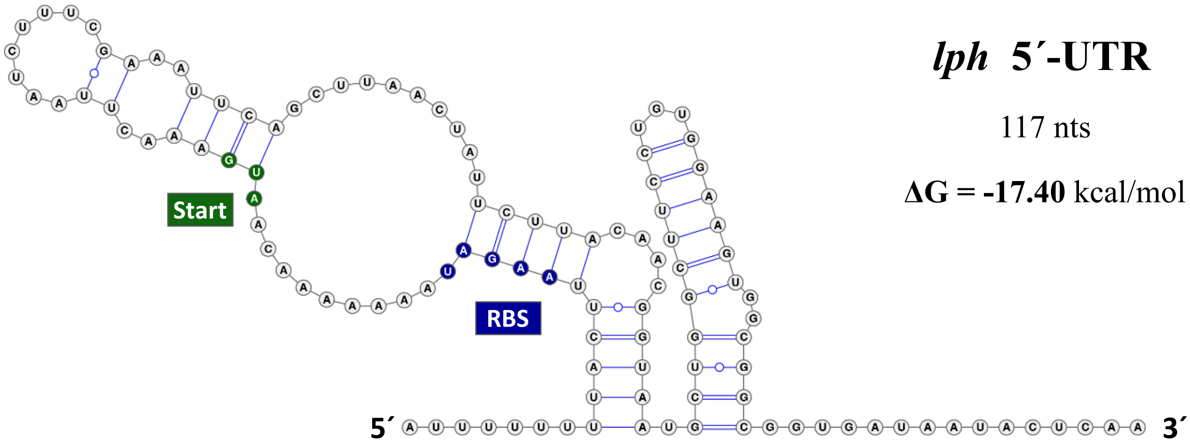


*lph* **5´-UTR**

117 nts

**ΔG = -17.40** kcal/mol

**S7 Fig.**

**S7 Fig.**

A) Promoters and 5´-UTR sequences of *pspC* in *S. pneumoniae.*

B) Promoters and 5´-UTR sequences of *lph* in *H. influenzae*

(A and B): promoter (green), transcriptional start site (arrows), ribosome binding site (RBS) (blue), coding region (shaded box).

C) Predicted RNA secondary structure of the 5´-UTR and 90 bases of *pspC*-mRNA.

D) Predicted RNA secondary structure of the 5´-UTR and 90 bases of *lph*-mRNA.

(C and D): RBS indicated in blue, start codon in green, start codon in green, putative secondary structure predicted with *RNAfold* and visualized with *VARNA* applet. Length of 5´-UTRs in nucleotides (nts) and ΔG values as indicated.

A

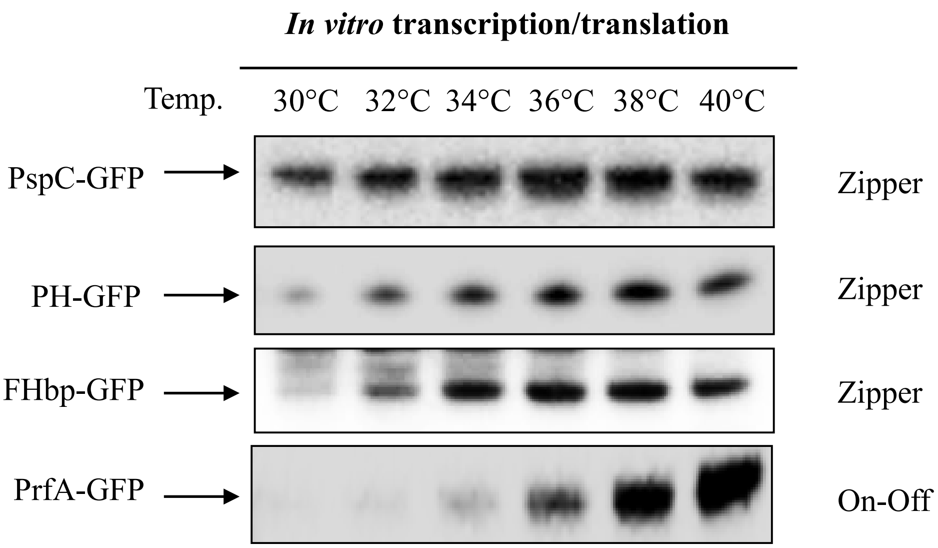
*In vitro* transcription/translation
Temp.
30°C
32°C
34°C
36°C
38°C
40°C
PspC-GFP
Zipper
PH-GFP
Zipper
FHbp-GFP
Zipper
PrfA-GFP
On-Off

S8 Fig.

**S8 Fig.**

*In vitro* transcription/translation assay comparing capsule producing enzymes with known thermosensors at a range of biological temperatures. PspC and PH show thermosensing. *N. meningitidis fHbp* and *L. monocytogenes* PrfA thermosensors are used as positive controls. Blots are representative of experiments performed on at least three occasions.
